# Increasing demand for plasma membrane contributes to the energetic cost of early zebrafish embryogenesis

**DOI:** 10.1101/775114

**Authors:** Jonathan Rodenfels, Pablo Sartori, Stefan Golfier, Kartikeya Nagendra, Karla Neugebauer, Jonathon Howard

## Abstract

How do early embryos apportion the resources stored in the sperm and egg? Recently, we established isothermal calorimetry (ITC) to measure heat dissipation by living zebrafish embryos and to estimate the energetics of specific developmental events. During the reductive cleavage divisions, the rate of heat dissipation increases from ∼60 nJ·s^−1^ at the 2-cell stage to ∼90 nJ·s^−1^ at the 1024-cell stage. Here we ask, which cellular process(es) drive these increasing energetic costs? We present evidence that the cost is due to the increase in the total surface area of all of the cells of the embryo. First, embryo volume stays constant during the cleavage stage, indicating that the increase is not due to growth. Second, the heat increase is blocked by nocodazole, which inhibits DNA replication, mitosis and cell division; this implicates some aspect of cell proliferation contributing to these costs. Third, the heat increase scales with total cell surface area rather than total cell number. Finally, the calculated costs of maintaining and assembling plasma membranes and associated proteins probably accounts for a significant proportion of the heat increase. Thus, the cell’s membrane is likely to contribute significantly to the total energy budget of the embryo.

**Highlight summary for TOC:** Rodenfels et al. measure the energetic costs of early zebrafish development, using calorimetry. Embryonic heat dissipation increases, but, more slowly than the number of cells during early cleavage stage development. Instead, the heat dissipation scales with the energetic cost associated with maintaining and producing new plasma membrane.

## Introduction

During early embryogenesis in oviparous animals, there is often a stage during which the cells divide without a change in total volume of the zygote (Tadros & Lipshitz, 2009). During this so-called reductive cleavage stage, components provided by the mother and stored in the oocyte, for example in the yolk, are used to build the early embryo. The constancy of volume allows one to study proliferative, developmental and metabolic processes independent of overall growth of the embryo. The embryo can be considered as an open system exchanging energy and matter with its environment. Thus, its metabolic and energetic profile that can be understood from basic principles, such as mass and energy conservation, stoichiometric constraints on production, homeostasis and morphometric parameter (Jusup, 2016). To investigate overall energetics, our lab has measured heat dissipation, which is equal to the net enthalpic change associated with all of the biochemical reactions taking place in the embryo (Rodenfels, 2019)

The heat dissipated during the development of frog (Nagano & Ode, 2014), fish (Rodenfels, 2019) and fly (Song, 2019) embryos can be measured using isothermal calorimetry (ITC). The measurements show, for example, that embryogenesis is exothermic, meaning that the medium surrounding the embryos heats up. In the zebrafish *Danio rerio*, cleavage stage lasts for ten divisions producing 1024 cells, referred to as the 1K stage. Recently, we showed that the heat dissipation oscillates in early zebrafish embryos, with an amplitude of ∼2% of the total heat dissipation (Rodenfels, 2019). The oscillations have a period equal to the cell cycle time and arise from heat dissipated by the biochemical reactions associated with the cell cycle oscillator, which coordinates events that take place during the cell division cycles. This analysis, based on non-invasive ITC measurements, shows the value of an approach that associates cellular events during development with energetics.

Concurrent with the oscillations, there is an overall increase in heat dissipation during the cleavage stage from ∼60 nJ·s^−1^ at the two-cell stage to ∼90nJ·s^−1^ at the 1K stage (Rodenfels, 2019). We call this the “increasing trend”. What cellular events account for the increasing trend in zebrafish embryos? In this work, we address this issue using a combination of pharmacological and computational approaches.

## Results

To ensure that the increasing trend is a robust phenomenon inherent to zebrafish embryos developing under of conditions, we measured enthalpic changes at three different temperatures that permit early embryogenesis while altering the duration of the cell cycle (Kimmel, 1995). Thirty embryos from one mother were manually synchronized to begin the second cleavage (time zero) within 3 minutes of each other and placed in an isothermal calorimeter (Figure 1A). At 28.5 °C, the heat dissipation rate increased from 60 ± 13 nJ·s^−1^ (mean ± standard deviation, *n* = 10 experiments each containing 30 embryos) to 88 ± 10 nJ·s^−1^ after 150 minutes, when the 10^th^ cleavage was complete (Figure 1B). At 23.5 °C, the initial dissipation rate of 51 ± 8 nJ·s^−1^ (*n* = 9 experiments) increased to 69 ± 10 nJ·s^−1^ after 210 minutes, when the 10th cleavage was complete (Figure 1C). At 33.5 °C, the initial dissipation rate of 82 ± 18 nJ·s^−1^ (*n* = 6 experiments) increased to 119 ± 22 nJ·s^−1^ after 130 minutes (Figure 1D). Thus, all three of the curves show a slow increasing trend during early embryogenesis, indicating its generality.

**Figure 1.**
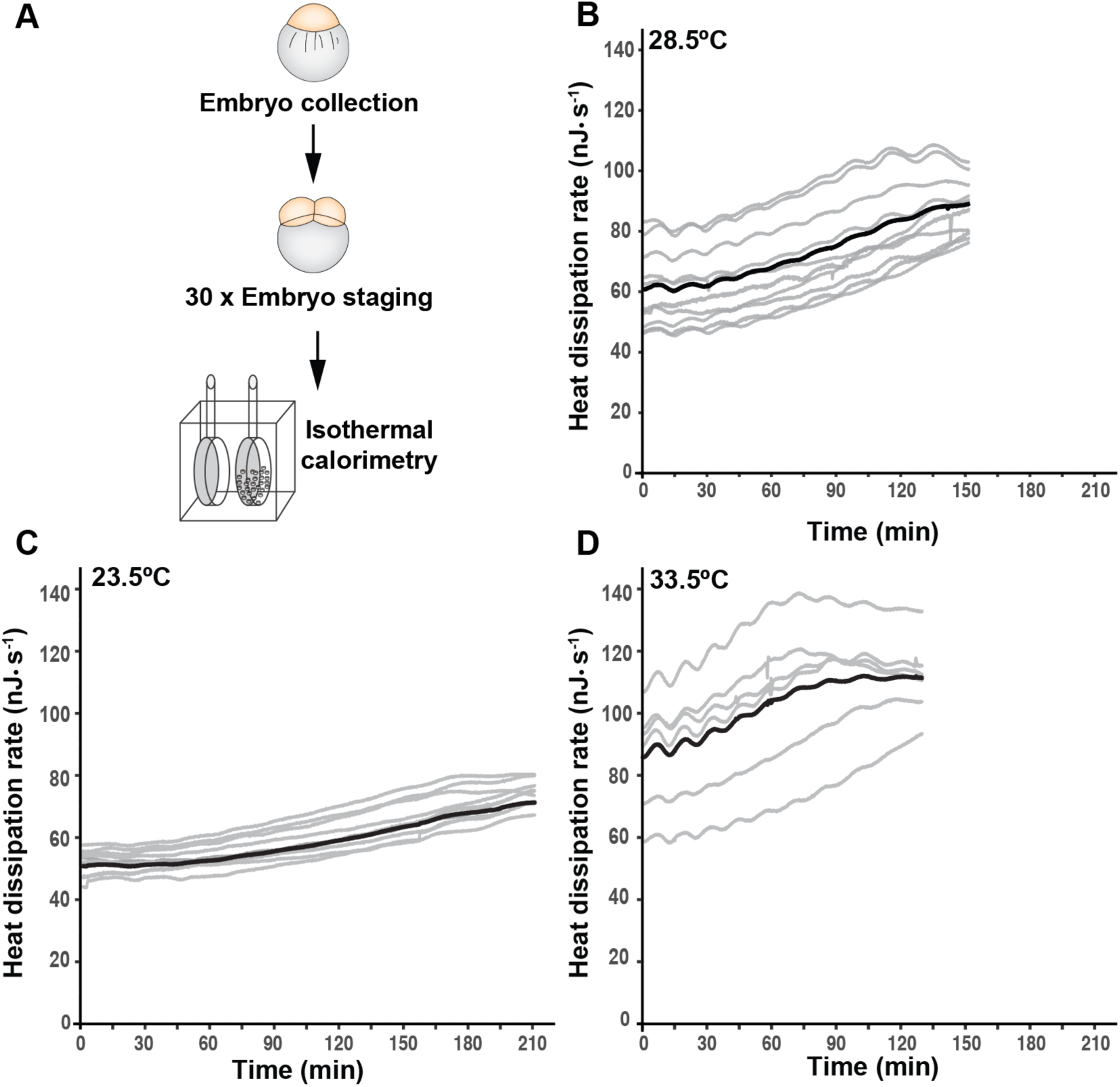
Heat dissipation in the early zebrafish embryo. **A** Schematic of an ITC experiment. Thirty embryos from a single pair of parents were collected and staged at the 2-cell stage. Following staging, the embryonic heat dissipation rate during development was measured using isothermal calorimetry. **B** The time-course of heat dissipation for 10 experiments at 28.5°C (gray lines) together with the mean (black line). Time zero corresponds to the beginning of cleavage at the 2-cell stage. Positive heat dissipation corresponds to heat transfer from the embryo to the surroundings. **C** The time-course of heat dissipation for nine experiments at 23.5°C (gray lines) together with the mean (black line). **D** The time-course of heat dissipation for 6 experiments at 33.5°C (gray lines) together with the mean (black line).

One possible explanation for the increasing trend is the increase in cell number. To test this possibility, we treated 2-cell embryos with nocodazole, which blocks microtubule assembly. As a consequence of microtubule impairment, nocodazole also blocks DNA replication, mitosis and cell division (Ikegami, 1997; Rodenfels, 2019). We showed previously that the embryos arrest development and the number of nuclei and cells remains at two throughout the first roughly 135 minutes of development, even though the small heat oscillations and the phosphorylation and dephosphorylation cycles associated with the cell cycle machinery continue (Rodenfels, 2019). Figure 2 shows that addition of nocodazole almost abolished the increasing trend: the relative increase in heat dissipation dropped from 46 ± 15 % in control (mean ± SD, *n* = 10) and 40 ± 23 % in DMSO treated cells (*n* = 6) to 13.5 ± 1.5 % in nocodazole-treated cells (*n* = 6). We conclude that some aspect of cell proliferation drives the increasing trend.

**Figure 2.**
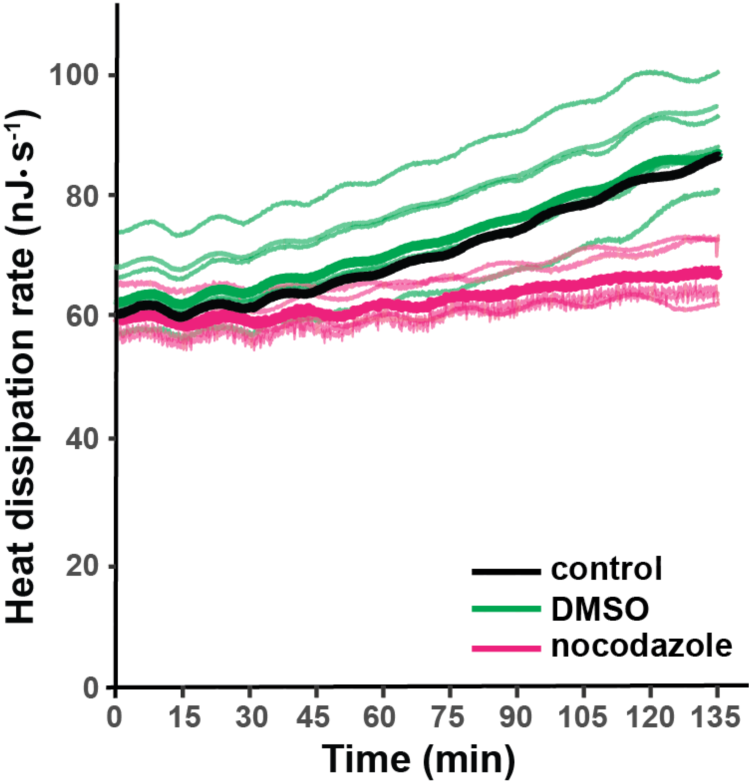
Nocodazole inhibits the increase in heat dissipation rate. Heat dissipation rate traces for 6 experiments in which nocodazole was added at the 2-cell stage. The thin magenta lines show individual traces and the thick magenta line is the mean. DMSO-treated control traces (n=6) are shown in thin green lines with the mean shown in the thick green line. The thick black line shows the mean trace from Figure 1B.

The results of nocodazole treatment suggest that the increasing trend is related to the number of nuclei or the number of cells. Therefore, we considered a model wherein the heat dissipation is proportional to cell volume [assumed to be constant] plus a term that is a function of the number of cells. This model is formalized by the following equation in which heat dissipation rate 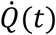 is a function of the number of cells (*N*) and therefore of the number of cell divisions (*n*):

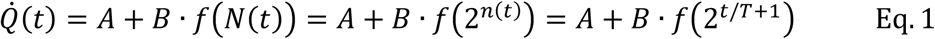

*A* is the constant volume component and *B* is a proportionality constant. *t* is time, *T* is the period of the cell cycle and *f* is a function (to be determined). Note that when *t* = 0 (the start of the experiment), *n* = 1 and *N* = 2.

If heat dissipation is proportional to the number of cells, then we expect that *f*(2^*t/T*+1^) ∝ 2^*t/T*^ because the number of cells increases exponentially while the doubling time, *T*, is approximately constant. If the heat increase is proportional to the rate of increase in the number of cells, then we expect *f*(*t*) ∝ (1*/T*)2^*t/T*^, which again shows a doubling time equal to the period of the cell cycle.

To test whether the heat scales with the number or increase in number of cells, we fit the heat increase with the following exponential equation:

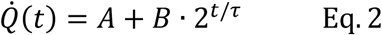

where *τ* is the heat doubling time. The equation was fit to the individual curves measured in each experiment. Comparison of the average fitted curve (using the average parameters from the individual fits) to the experimental traces at 28.5 °C shows that the model accords with the data (Figure 3A). The data from the ten individual experiments (*Q*_*i*_(*t*), *i* = 1, …, 10) were rescaled by subtracting *A*_*i*_, dividing by *B*_*i*_ and plotting against time divided by *τ*_*i*_. The rescaled curves showed a “data collapse” (Bhattacharjee & Seno, 2001), indicating that the model in Equation 2 is a good description of the data, though it fits less well during the last two cell divisions (Figure 3B). The curves were approximately linear when the rescaled data were plotted on a semi-log plot, indicating that the increase is exponential (Figure 3B, inset). The mean and standard deviations of *A, B* and *τ* from the individual fits are compiled in Table 1, together with the mean and standard deviation of the cell cycle periods *T*. Thus, Equation 2 is a good empirical description of the increasing trend during cleavage stage.

**Table 1:** Means and SDs of the parameters in surface-are model. *A, B*, and *τ* in Equation 2 were fit to the individual experimental curves at 28.5 °*C* (Figure 1A, *n* = 10), 23.5 °*C* (Supplementary Fig. 1A, *n* = 9) and 33.5 °*C* (Supplementary Fig. 1B, *n* = 6). The mean cell doubling time, *T*, is equal to the mean oscillatory period.

| Parameter | Mean ± SD | | |
| --- | --- | --- | --- |
| Temperature | 28.5°C | 23.5°C | 33.5°C |
| Volume term, $A$ | $52 \pm 12 \text{ nJ} \cdot \text{s}^{-1}$ | $47 \pm 1.8 \text{ nJ} \cdot \text{s}^{-1}$ | $88.4 \pm 22.4 \text{ nJ} \cdot \text{s}^{-1}$ |
| Area prefactor, $B$ | $8.2 \pm 3.2 \text{ nJ} \cdot \text{s}^{-1}$ | $5.12 \pm 2.9 \text{ nJ} \cdot \text{s}^{-1}$ | $4.22 \pm 0.9 \text{ nJ} \cdot \text{s}^{-1}$ |
| Heat doubling time, $\tau$ | $60.4 \pm 10.4 \text{ min}$ | $74.3 \pm 14.9 \text{ min}$ | not well fit |
| Cell doubling time, $T$ | $17.2 \pm 0.8 \text{ min}$ | $23.4 \pm 1.4 \text{ min}$ | $14.2 \pm 0.8 \text{ min}$ |

**Figure 3.**
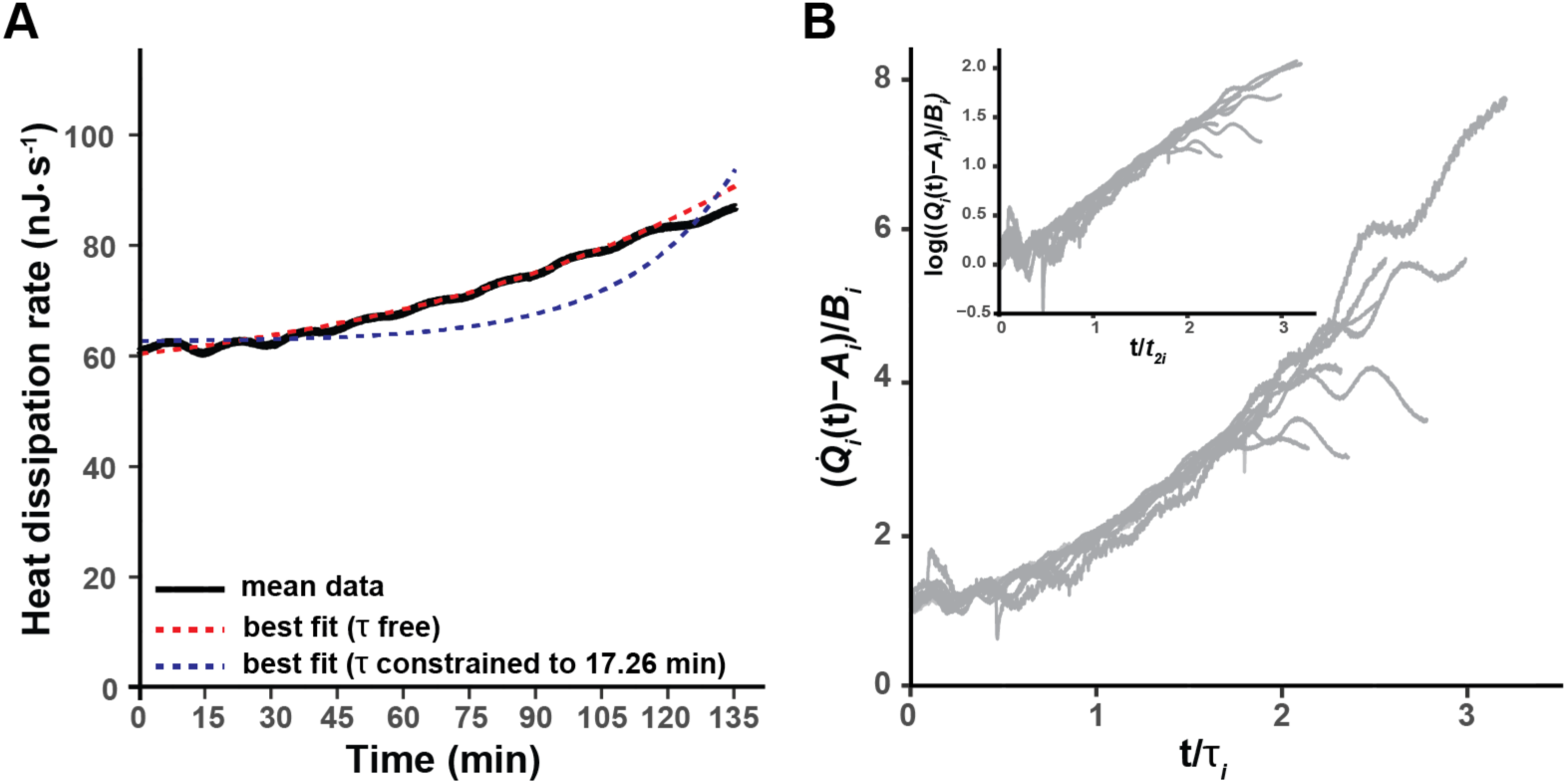
The heat dissipation increase accords with a slow exponential with half time approximately three times the cell cycle period. **A** Least-squares fit of the heat dissipation curves to an exponential plus a constant,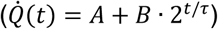. *A, B* and *τ* are free parameters. The fits were done on the individual experimental curves (gray lines in Figure 1) and averaged (the red dotted line). The mean of the experimental heat dissipation curves is shown in black. The blue dotted curve is the least-squares fit with *τ* constrained to be equal the average cell cycle time of 17.3 min with *A* and *B* free parameters. **B** The heat dissipation trajectories from the ten individual experiments (*Q*_*i*_(*t*), *i* = 1, …, 10) were rescaled by subtracting *A*_*i*_ and dividing by *B*_*i*_ plotted against time divided by *τ*_*i*_. The superimposed curves illustrate the exponential rise, which is also apparent in the linear increase on a log-linear scale (Inset).

We can now ask how the doubling time of the increasing trend relates to the doubling time of the cells. Interestingly, the doubling time of the increasing trend, *τ* = 60 ± 10 min (mean ± SD, *n* = 10), was about three times longer than the average period of the cell divisions (*T* =17.2 ± 0.8 min). In other words, heat dissipation increased considerably more slowly than expected if the heat were proportional to the number of cells or the rate of increase in the number of cells. Figure 3A (blue curve) also shows a theoretical curve with *τ* being constrained to the cell division time of 17.2 min. We conclude from this poor fit that the data are not compatible with the heat dissipation rate doubling every cell cycle.

The increasing trend was also well fit by Equation 2 at lower temperatures. At 23.5 °C the doubling time of the increasing trend, *τ* = 74 ± 15 min, was again about three times longer than the average period of the cleavage divisions (*T* =23 ± 2 min). Comparison of the average fit curve to the average of the experimental traces shows the model accords with the data at lower temperatures (Table 1, Supplementary Fig. 1A). The heat dissipation curves at 33.5 °C were more variable from experiment to experiment, and some curves showed decreasing heat dissipation at the end of cleavage stage development, raising the possibility that the health of the embryos is compromised at high temperatures. Fitting Equation 2 to the average of the individual traces gave a heat doubling time of *τ* = 68 min, which is about 5 times larger than the average cell cycle time (*T* =14.2 ± 0.8 min) at 33.5 °C. The fit was not good, but significantly better than when *τ* was constrained to the average cell doubling time, i.e. *τ* = *T* (Supplementary Fig. 1B). Thus, at all temperatures, the increasing trend was much slower than expected if it scaled with cell number. Taken together, these findings are inconsistent with the increasing trend being to an exponential increase in the numbers of cells, genomes or nuclei.

An alternative hypothesis is that the slower doubling time is due to a heat dissipation rate proportional to total cell surface area. To explore this possibility, we derived a simple model that shows that the total cell surface area, as well as the change in total cell surface area, doubles three times more slowly than the number of cells (Supplementary Material). This cell surface model assumes that (i) the cells are spherical; (ii) the total volume of the cells is constant throughout the cleavage stage (the initial volume is 60 × 10^6^ μm^3^); and (iii) the cell doubling time is constant throughout the cleavage stage. We defer consideration of potential deviations from the assumptions to the Discussion section. With this caveat, our data agree with the predictions of the cell surface model (Table 1).

To further test the cell surface model, we asked whether the energetic costs associated with the increasing trend are within the range of values, derived from the literature, for maintaining and/or producing plasma membrane. To make this comparison, we write the proportionality constant *B* in Equation 2, as

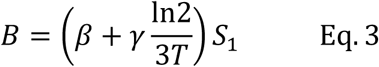

(Supplementary Material), where *β* is the maintenance cost per unit area and unit time (nJ·s^−1^·μm^−2^) and *γ* is the production cost per unit area of building new membrane (nJ·μm^−2^). *S*_1_ is the total area (μm^2^) at the two-cell stage, assuming spherical cells, and *T* is the average cell doubling time.

The cost of maintaining surface area includes the work done by ion pumps to maintain the membrane potential against leakage through ion channels and transporters, work done by flippases and floppases to maintain the asymmetry of the lipids, and work done by fission and fusion machinery such as the GTPase dynamin and the ATPase NSF associated with vesicular endocytosis and exocytosis. To estimate the total maintenance cost, we assumed a total protein density of 40 · 10^3^ *μm*^−2^ (Itzhak, 2016, Quinn, 1984) that 3% are ATPases and that the average ATPase activity is 3 s^−1^ Membrane protein turnover is another maintenance cost, which we estimate assuming an average protein half-life of 42h (Peshkin, 2015), protein degradation by lysosomes, and re-synthesis by the ribosome. Assuming that the hydrolysis of ATP generates 40 kJ·mol^−1^ of enthalpy, the energy cost of maintaining the plasma membrane ATPase activity is *β*_ATPase_ = 0.24 fJ·s^−1^·μm^−2^ and the cost of plasma membrane protein turnover is *β*_turnover_ = 0.02 fJ·s^−1^·μm^−2^ (Table 2). Inserting *β* = *β*_ATPase_ + *β*_turnover_ into Equation 3 results in a total maintenance heat dissipation of 0.24 nJ·s^−1^ for a 2-cell embryo with an initial surface area of *S*_1_ = 930 × 10^3^ μm^2^.

**Table 2:** Estimated energetic parameters at 28.5 °C.

| Parameter | Estimates |  | Values from fits |
| --- | --- | --- | --- |
| Volume term<sup>1</sup>, *A* | 90 nJ · s<sup>-1</sup> |  | 52 ± 12 nJ · s<sup>-1</sup> |
| Area pre-factor<sup>2</sup>, *B* | $\beta$ (maintenance) | $\gamma$ (production) | |
| | $\beta_{\text{ATPase}}^3 = 0.24 \cdot 10^{-3}$<br>pJ · s<sup>-1</sup> · $\mu\text{m}^{-2}$ | $\gamma_{\text{lipid}}^4 = 0.12 \text{ to } 24 *$<br>pJ · $\mu\text{m}^{-2}$ | |
| | $\beta_{\text{turnover}}^5 = 0.02 \cdot 10^{-3}$<br>pJ · s<sup>-1</sup> · $\mu\text{m}^{-2}$ | $\gamma_{\text{protein}}^6 = 4.3 \text{ to } 128 *$<br>pJ · $\mu\text{m}^{-2}$ | |
| | $B = \left( \sum \beta_i + \frac{\ln(2)}{3T} \sum \gamma_i \right) \cdot S_1 = 0.9 \text{ to } 32 \text{ nJ} \cdot \text{s}^{-1} *$ (see<sup>7,8</sup>) | | 8.2 ± 3.2 nJ · s<sup>-1</sup> |

<sup>1</sup> $A = 90 \text{ nJ/s}$ . From (Lynch & Marinov, 2015) based on the volume of $60 \cdot 10^6 \mu\text{m}^3$ and a temperature of 28.5°C
<sup>2</sup> $B = \left( \beta + \gamma \frac{\ln 2}{3T} \right) S_1$
<sup>3</sup> $\beta_{\text{ATPases}} = 0.24 \cdot 10^{-3} \text{ pJ} \cdot \text{s}^{-1} \cdot \mu\text{m}^{-2}$ . We assume that the density of proteins is $40 \cdot 10^3 \mu\text{m}^{-2}$ , that 3% are ATPases and that the average ATPase activity is 3/s. $\Delta H_{\text{ATP}} = 40 \text{ kJ} \cdot \text{mol}^{-1}$
<sup>4</sup> $\gamma_{\text{lipid}} = 0.12 - 24 \text{ pJ} \cdot \mu\text{m}^{-2}$ . *lower bound*: We assume that there are $0.9 \cdot 10^6 \text{ molecules} \cdot \mu\text{m}^{-2}$ of phospholipids and $0.6 \cdot 10^6 \cdot \mu\text{m}^{-2}$ molecules cholesterol in the plasma membrane. From (Lynch & Elife, 2017), we assume that the phospholipid cost is just adding the head groups, which is 2 ATP equivalents per phospholipid and zero for the hydrolysis of stored cholesteryl-esters. This ignores moving lipids and cholesterol from yolk granules in the yolk to the membrane bilayer (and flipping across the bilayer) and also any possible synthesis or head groups. $\Delta H_{\text{ATP}} = 40 \text{ kJ} \cdot \text{mol}^{-1}$ . *Upper bound*: Takes synthesis of acyl chains, head groups and cholesterol into account and is 200x times due to increased ATP cost for de-synthesis (Lynch & Elife, 2017).
<sup>5</sup> $\beta_{\text{turnover}} = 0.02 \cdot 10^{-3} \text{ pJ} \cdot \text{s}^{-1} \cdot \mu\text{m}^{-2}$ . We assume that the density of proteins is $40 \cdot 10^3 \mu\text{m}^{-2}$ with an average half-life time of 42h from (Peshkin et al., 2015). We assume the cost of protein degradation by the lysosome is zero ATP. We account for endocytic GTPase activity in the ATPase term. The cost of re-synthesis of degraded proteins is 4 ATP/aa and the average membrane protein is 400 aa. $\Delta H_{\text{ATP}} = 40 \text{ kJ} \cdot \text{mol}^{-1}$
<sup>6</sup> $\gamma_{\text{protein}} = 4 - 128 \text{ pJ}/\mu\text{m}^2$ . *lower bound*: From (Lynch & Marinov, 2015) we assume that the density of proteins is $40 \cdot 10^3 \mu\text{m}^{-2}$ . Then if there is no synthesis and we ignore putting them into the plasma membrane, then the cost is 4 ATP/aa. We assume the average protein is 400 aa. $\Delta H_{\text{ATP}} = 40 \text{ kJ} \cdot \text{mol}^{-1}$ . *Upper bound*: Full synthesis is 30 times more because the average ATP cost for synthesizing an amino acid is 30 ATP (Lynch & Marinov, 2015)
<sup>7</sup> $B = \left( \beta_{\text{ATPase}} + \beta_{\text{turnover}} + (\gamma_{\text{lipid}} + \gamma_{\text{protein}}) \frac{\ln 2}{3T} \right) S_1 = 0.9 \text{ nJ/s}$ (lower bound) and $32.0 \text{ nJ/s}$ (upper bound)
<sup>8</sup> $S_1 = 930 \cdot 10^3 \mu\text{m}^2$ ( $S_1 = S_1 \cdot 0.7$ for $\gamma_{\text{lipid}}$ ). The area at the two-cell stage, assuming spherical cells and a total volume of $60 \cdot 10^6 \mu\text{m}^3$
\* The lower estimates are assuming synthesis from pre-assembled acyl chains, lipid headgroups and amino acids. The higher estimate corresponds to *de novo* synthesis of the building blocks.

The cost of producing new plasma membrane includes synthesizing both lipids and proteins. The one-cell embryo has high levels of maternally loaded phospholipids, cholesterol and cholesteryl-esters; *de novo* synthesis of membrane lipids is therefore probably absent during cleavage stage development in zebrafish (Fraher, 2016). Even if new phospholipids need to be assembled from diacyl chains derived from triglycerides in lipoprotein-like yolk granules (i.e. no synthesis of fatty acids) and preexisting head groups, the cost is quite modest: the removal of one fatty acid and the addition of the head group is estimated to utilize 2 CTPs (Lynch, 2017), corresponding to 2 ATP-equivalents per lipid. The number of phospholipids and cholesterol molecules is calculated from the lipid area (0.65 nm^2^), assuming that 70% of the membrane is occupied by 60% phospholipid and 40% cholesterol. This results in a lipid production cost of 0.12 pJ·μm^−2^ (*γ*_lipid_,Table 2). To estimate the production cost of membrane proteins, we assume a density of 40 × 10^3^ μm^−2^, an average size of 400 amino acids, and 4 ATP per amino acid. Assembly of plasma membrane proteins costs 4.3 pJ·μm^−2^ (*γ*_protein_,Table 2). This results in a heat dissipation from production costs of 0.9 nJ·s^−1^ for a 2-cell embryo (using an initial surface area of *S*_1_ = 930 × 10^3^ μm^2^ and an average cell doubling time of 17.2 min). If phospholipids, cholesterol or amino acids need to be synthesized *de novo*, then the costs can be 50 to 200 times more due to the substantial increase in ATP hydrolysis associated with the *de novo* synthesis of fatty acids, amino acids and cholesterol.

We now compare the measured values of *A* and *B* with the estimated values from the literature. The parameter *A* corresponds to the energetic cost of maintaining the fixed volume of cytoplasm in the early embryo. We assume that this is equal to the basal maintenance cost of cells and organisms (Lynch & Marinov, 2015), which is related to the basal metabolic rate (Makarieva et al., 2008). The basal maintenance cost has been estimated as a function of volume over a wide range of cell and organism volumes that encompass the volume of the zebrafish embryo. Using the value from (Lynch & Marinov, 2015) normalized to 28.5 °C and 23.5 °C, we obtained an estimated value of *A* to be 90 nJ·s^−1^ at 28.5 °C and 65 nJ·s^−1^ at 23.5 °C. This is within a factor of two of the value measured from the curve fits of 52 nJ·s^−1^ at 28.5 °C and 47 nJ·s^−1^ at 23.5 °C (Table 1). Thus, the initial rate of heat dissipation, which is mainly determined by the parameter *A* in our model, is well accounted for by the typical energetic demands of cells.

The estimated lower bound value of *B* from the literature, which includes contributions from both maintenance and production of surface area, is 0.92 nJ·s^−1^ (*B*, Table 2), assuming that there is no *de novo* protein or lipid synthesis. This value is roughly an order of magnitude smaller than the value obtained from our curve fits, 8.2 ± 3.4 nJ·s^−1^. If we include fatty acid and/or amino acid synthesis, then the estimated costs of producing plasma membrane are much higher, and the upper bound for *B* would be 32 nJ·s^−1^. (Table 2). Thus, if all lipids or proteins were synthesized *de novo*, the value for B would exceed the measured value. If, however, ∼25% of the required plasma membrane lipids and proteins were to be synthesized *de novo*, our estimate for *B* would be equal to the value of *B* obtained from our curve fits.

## Discussion

In this work, we have investigated the overall energetics of cell proliferation without growth during embryogenesis, taking advantage of reductive cleavage divisions as a model system. Our experimental measurements in the early zebrafish embryo show that the rate of heat dissipation increases from an initial value of ∼60 nJ·s^−1^ at the two-cell stage to ∼90 nJ·s^−1^ (at 28.5 °C) at completion of the tenth division. This suggests an energetic cost of proliferation in the embryo, leading us to consider several hypotheses for the source of this cost. Because the volume of the embryo is constant, we ruled out growth and instead used experimental and computational approaches to evaluate the possibilities that the increase in energetic costs may be due to DNA replication, mitosis, the production of new plasma membranes or any other process taking place in the cytoplasm or nucleus. Based on our findings, we argue that the cost is likely due to the increase in the total surface area of all of the cells of the embryo, i.e. the production and/or maintenance of plasma membranes that enclose the increasing number of cells of the embryo. We discuss the key experiments below.

The crucial finding that specifically implicates cell proliferation is that the heat increase is blocked by nocodazole. This drug inhibits DNA replication, mitosis and cell division but does not kill the embryo. If the heat increase were still measurable after treatment, then some other process within the cytoplasm e.g. a hypothetical expansion of the number of organelles, could account for the heat increase. This is not the case, implicating some aspect of cell proliferation contributing to these costs. An additional argument consistent with this interpretation is provided by our experiments testing the effects of temperature. We show that the rate of the increase in heat dissipation correlates with the length of the cell cycle, which slows at lower temperatures and speeds up at higher ones. These data also confirm the robustness of the increase in energetic costs to different environmental conditions.

The second key finding is computational. We show that the heat increase scales with total cell surface area rather than total cell number. The increase in the dissipation rate follows a slow exponential with a doubling time approximately three times longer than the doubling time of the cells. The slow doubling time is inconsistent with increased cost due to increased DNA, chromatin, nuclear volume and nuclear area, which all correlate with the number of cells in fish and frog (Gerhart, 1980, Keller, 2008)).

The slow doubling time is consistent with the surface-area model in which the heat dissipation is proportional to total cell area and to the rate of change of total cell area. To implement this model, we made a number of assumptions that need to be discussed. One assumption is that the cells are spherical. However, at the 2-cell stage, the two cells appear hemispherical and are open at their bases. Therefore, we may be overestimating the initial surface area at the 2-cell stage. Because the surface area contribution is small at early stages, this underestimate is not likely to have a large impact on the increasing trend. Another assumption is the embryo’s constant volume. Cellularization is not complete until the 64-cell stage (Kane & Kimmel, 1993; Karlstrom & Kane, 1996), after which the constant volume assumption is well accepted. Before the 64-cell stage, we may be underestimating the cytoplasmic volume contributing to the embryo’s energetics. A third assumption is that the cell cycle time is constant. However, the cell cycle slows down by ∼30% over the last three cell divisions. This will dilate the time axis towards the end of the cleavage stage. We have also included this in Supplementary Materials where we refitted the heat dissipation curves with a modified model where *T* is now a function of time. The curve fits to the mean trend heat dissipation curve are the shown in Supplementary Figure 2. Correcting for the change in period over time does not improve or worsen the curve-fitting, indicating that a constant cell cycle time is a reasonable assumption. In summary, there may be deviations from our assumptions that could in principle influence the kinetics of the increasing trend, especially at early stages before cellularization. However, we have argued that these effects must be modest and are not likely to alter our conclusions.

Additional calculations provide validation of our working model that increasing energetic demands associated with plasma membrane increase heat dissipation. Based on literature values of maintenance costs of membrane ATPases, and the assembly costs of lipids and membrane protein (from premade lipid and amino-acid components), the maintenance and addition of surface area in the form of plasma membranes can account for roughly 10% of the measured increase. If *de novo* synthesis is required for 25% of the phospholipids, cholesterol and proteins, then up to 100% of the measured increase might be accounted for. Thus, the calculated costs of maintaining and assembling plasma membranes and associated proteins likely accounts for a significant proportion of the heat increase. Our work motivates future studies that will investigate the sources of precursors, the costs of their biosynthesis, and the real fluxes of biomolecules among different cellular compartments.

Is the increasing energetic cost of cell proliferation conserved to other embryonic organisms? And why does heat dissipation level off after cleavage stage? Although *Xenopus leavis* exhibits holoblastic cleavage divisions (meaning the yolk is contained within each of the cells), a similar slow exponential increase in heat dissipation has been observed (Nagano & Ode, 2014). Interestingly, the increase in heat dissipation slows down after cleavage and again after gastrulation during frog development (Nagano & Ode, 2014). These slow-downs coincide with the embryos mid-blastula transition and gastrulation during which the embryonic cell cycle lengthens by about 3-fold and 8-fold respectively (Ferree, 2016; Siefert, 2015). A longer cell cycle time and, thus, a slower increase in surface area, could account for this observed slow-down. The presence of G1 and G2 cell cycle phases may also suggest that embryonic cells transition to grow in volume and thus are burden by the energetic cost of synthesizing new new cytoplasm.

## Materials and Methods

### Zebrafish husbandry and staging

Adult zebrafish were maintained and bred under standard conditions. Wild-type (AB) embryos were left to develop in E3 medium (5 mM NaCl, 0.17 mM KCl, 0.33 mM CaCl_2_, mM MgSO_4_) to the desired stage. The temperature was 28.5°C unless otherwise indicated. Pairs of fish were paired for a maximum of ten minutes after which eggs were collected and allowed to develop for 30 minutes at 28.5°C. Staging was done based on morphology. 30 embryos were selected such that their first cleavage furrows initiated within 3 minutes of each other. In this way, the population of embryos was synchronized at the 2-cell stage.

### Isothermal calorimetry (ITC)

Calorimetry experiments were carried out using a Malvern MicroCal VP-ITC (Malvern Instruments Ltd, Worcestershire, UK). The temperature in the instrument was set to desired temperatures of 22.5 °C, 28.5°C and 33.5 °C. The reference power (μcal s^−1^) was set to 11.5, the initial injection delay after calibration was set at 240s, the feedback mode/gain was set to high, and the ITC equilibration options were set to “fast equilibration & auto”. The injection syringe parameters were set as follows: injection volume, 2 μl; duration, 2 seconds; spacing, 14,400 s (240 min); number of injections, 3. The ITC experiments were performed without the injection syringe and stirring and the ITC chambers were covered with a plastic lid. The sample cell was filled with either 1.57 ml of E3 medium or E3 medium with the desired concentration of chemical inhibitors. The reference cell was filled with water.

### Data analysis and curve fitting

Data analysis was performed in R (version 3.5.3) using RStudio (version 1.2.1335) with the additional libraries “`ggplot2`”, “`tidyr`”, “`stats`”, and “`dplyr`”. Each experiment, corresponding to a biological replicate, is an ITC measurement of a group of 30 staged zebrafish embryos; n represents the number of experiments. Statistical parameters including the exact value of n are reported in the text or figure/table legends Curve fitting was performed by nonlinear least-squares fits of Equation 2 to single heat dissipation trajectories (groups of 30 embryos) using the `nls()` function in R.

## ACKNOWLEDGMENTS

We thank the participants of the 2019 Physical Biology of the Cell course at the Marine Biology Laboratory in Woods Hole, MA for helpful discussion during the inception of this project. This work was supported by funding from an EMBO Long-term Fellowship ALTF 754-2015 (to JR), the Eric and Wendy Schmidt Membership in Biology at the Institute for Advanced Study (to PS), NIH R21 HD094013 (to KMN) and NIH R01 GM110386 (to JH). Its contents are solely the responsibility of the authors and do not necessarily represent the official views of the NIH.

---

**Supplementary Figure 1.**
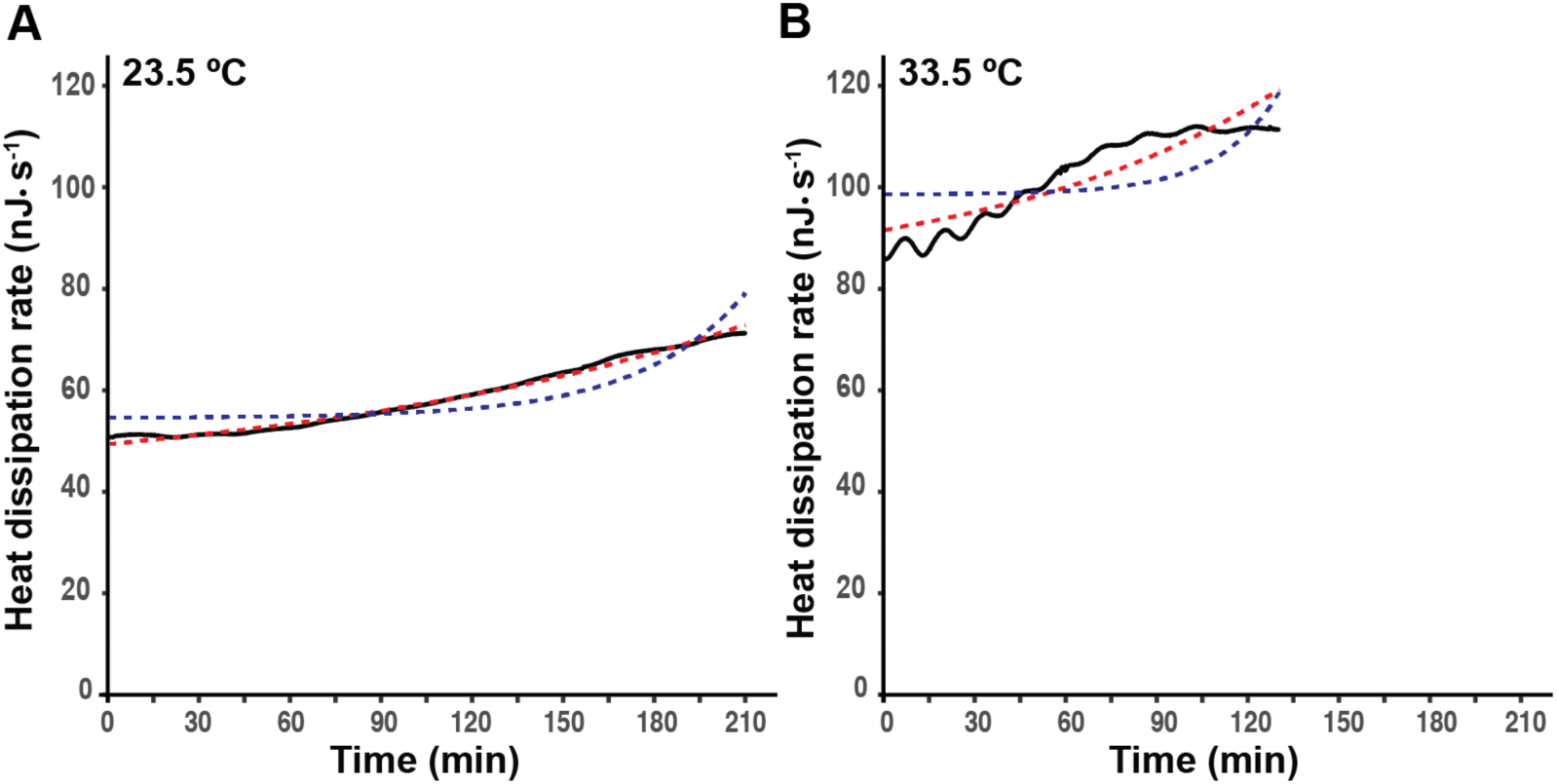
The heat dissipation increase accords with a slow exponential with half time approximately three times the cell cycle period at different temperatures. **A** Least-squares fit of the heat dissipation curves at 23.5 °C to an exponential plus a constant, 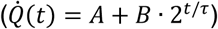. *A, B* and *τ* are free parameters. The fits were done on the individual experimental curves (gray lines in Figure 1C) and averaged (the red dotted line). The mean of the experimental heat dissipation curves is shown in black. The blue dotted curve is the least-squares fit with *t*_2_ constrained to be equal the average cell cycle time of 24.2 min with *A* and *B* free parameters. **B** Least-squares fit of the heat dissipation curves at 33.5 °C to an exponential plus a constant,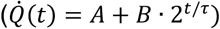. *A, B* and *t*_2_ are free parameters. The fits were done on the mean experimental curve. The mean of the experimental heat dissipation curves is shown in black and the fit is shown as the red dotted curve. The blue dotted curve is the least-squares fit with *τ* constrained to be equal the average cell cycle time of 14.2 min with *A* and *B* free parameters.

**Supplementary Figure 2.**
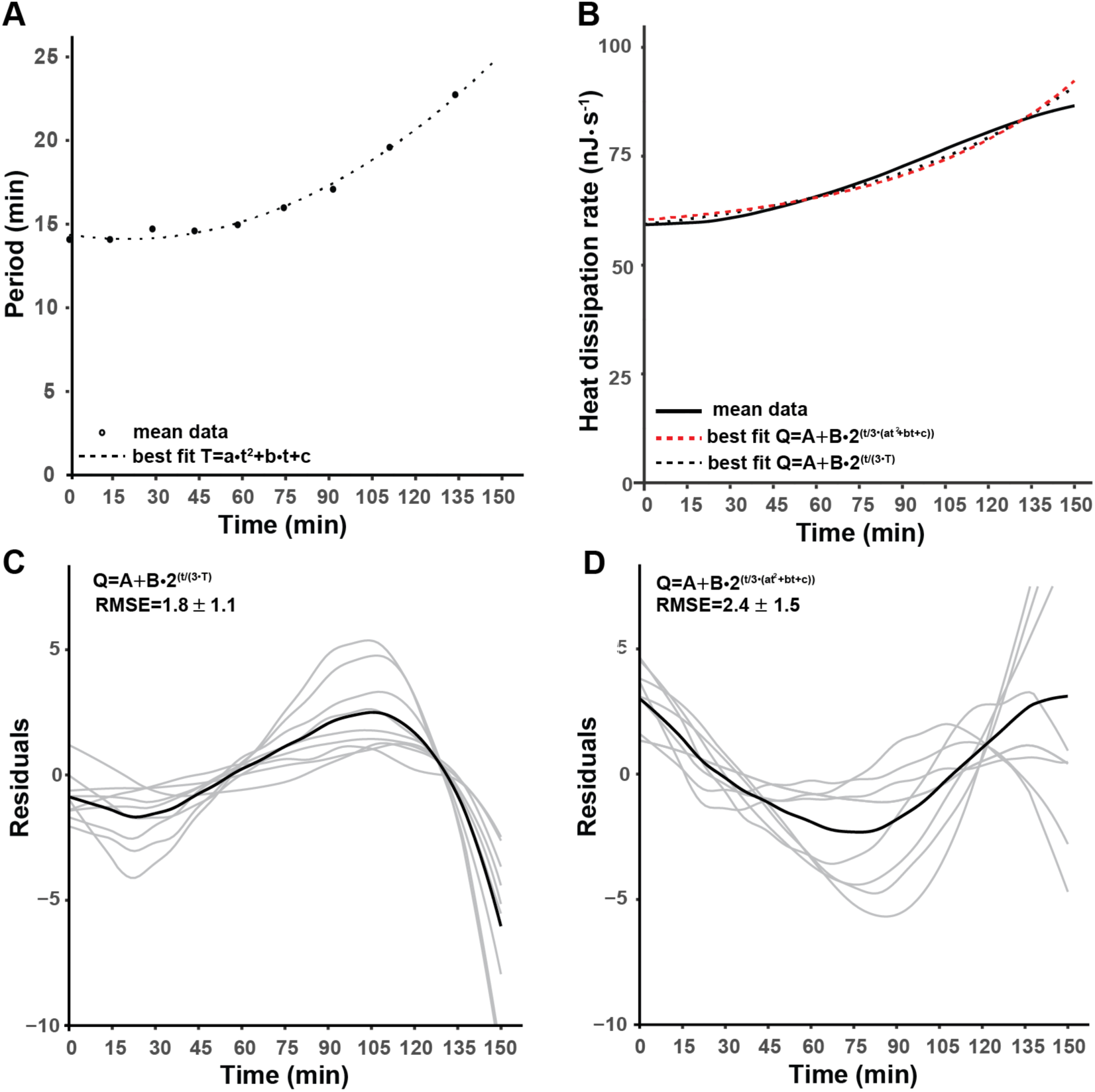
Accounting for an increase in cell cycle time. **A** Least-squares fit of the mean periods at 28.5 °C to *T*(*t*) = *at*^2^ + *bt* + *c*, black dotted line where *a, b* and *c* are free parameters. **B** Least-squares fits of the mean trend heat dissipation curve at 28.5 °C to 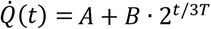, black dotted line and 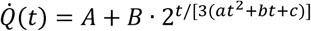, red dotted line, where *A, B* free parameters and *a, b* and *c* where constrained by the best fits of the individual experiments by the equation shown in A. **C & D** Residuals of the fits of the least square fits show in B to individual experiments, gray lines and their average (black line). Comparison of the root mean square error (RMSE) of the residuals of the individual experiments shows that neither model fits better or worse.

## Supplementary Materials

### Surface-area model

The surface-area model assumes that

(i) The cells are closed spheres.

(ii) The total volume (60 · 10^6^ μm^3^) remains constant over the cleavage stage

(iii) Cell division time is constant

The parameters in the model are defined as follows:

| Parameter | Meaning | Value |
| --- | --- | --- |
| $\dot{Q}(t)$ | Heat dissipation rate | 60-85 nJ·s<sup>-1</sup> |
| $t$ | Time from the beginning of cell 2 cleavage | 0 – 150 minutes |
| $n = t/T + 1$ | Number of divisions, $T$ is the division period | 1-10 |
| $N_n = 2^n$ | Number of cells* | 2-1024 |
| $R_n$ | Radius after the $n^{\text{th}}$ division | $R_0 = 243 \mu\text{m}$ |
| $S_n$ | Total surface area after the $n^{\text{th}}$ division | $S_1 = 930 \cdot 10^3 \mu\text{m}^2$ |
| $V_n$ | Total volume after the after the $n^{\text{th}}$ division | $V_0 = 60 \cdot 10^6 \mu\text{m}^3$ |

When *n* = 0 (i.e. the number of cells *N* = 2^0^ = 1), the volume and surface are:

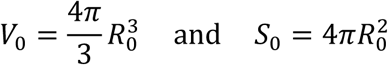

Because *V*_0_ = 60 · 10^6^ μm^3^, *R*_0_ = 243 μm.

After *n* divisions, the total cell volume is

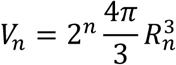

Because total volume is constant, *V*_5_ = *V*_0_, and so

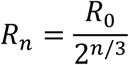

Thus,

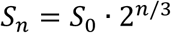

In other words, the total surface area of all the cells in the embryo double every third cell division.

To write the area as a function of time, rather than the number of cell divisions, we note that *n* = *t/T* + 1, where *T* is the doubling time. This formula correctly predicts that at the two-cell stage, *t* = 0, *n* = 1 and *N* = 2. Thus, in continuous time,

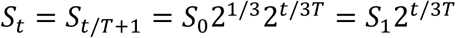

where *S*_1_ = 930 · 10^3^ μm^2^ is the area at the two-cell stage.

Note that differentiation of this last equation shows that the change in surface area *dS*/*dt* (*t*) is also proportional to 2^*t/*3*T*^. Thus, the time-dependence of the increase in heat dissipation is consistent with it being due to processes proportional to the total cell surface area or to the change in surface area.

Substituting this last equation into Equation 1 from the main text, we obtain

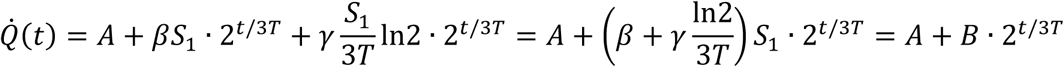

where *A* is a constant term (e.g. proportional to volume), the second term is proportional to surface area, and the third term is proportional to the increase in surface area.

### Correction of the model for the slowing of the cell cycle

We assume that the change in period over time has the functional form:

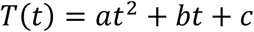

We obtained *a, b* and *c* for each biological replicate by curve-fitting and modified our heat dissipation model the following:

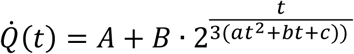

Where *A, B* are free parameters. The obtained mean values for *A* = 58 ± 14 nJ · s^−1^ and *B* = 6.8 ± 2.9 nJ · s^−1^are statically not different from obtained values from the original model (see Table 1), *p* = 0.288 and *p* = 0.299, students *t*-test.

